# Glial response to hypoxia in *trachealess* mutants induces synapse remodeling

**DOI:** 10.1101/527671

**Authors:** Pei-Yi Chen, Yi-Wei Tsai, Angela Giangrande, Cheng-Ting Chien

## Abstract

Synaptic structure and activity are sensitive to environmental alterations. Modulation of synaptic morphology and function is often induced by signals from glia. However, the process by which glia mediate synaptic responses to environmental perturbations such as hypoxia remains unknown. Here, we report that, in the *Drosophila trachealess* (*trh*) mutant, smaller synaptic boutons form clusters named bunch boutons appear at larval neuromuscular junctions (NMJs), which is induced by the reduction of internal oxygen levels due to defective tracheal branches. Thus, the bunch bouton phenotype in the *trh* mutant is suppressed by hyperoxia, and recapitulated in wild-type larvae raised under hypoxia. We further show that hypoxia-inducible factor (HIF)-1α/Similar (Sima) is critical in mediating hypoxia-induced bunch bouton formation. Sima upregulates the level of the Wnt/Wingless (Wg) signal in glia, leading to reorganized microtubule structures within presynaptic sites. Finally, hypoxia-induced bunch boutons maintain normal synaptic transmission at the NMJs, which is crucial for coordinated larval locomotion.

**Author summary:** Oxygen is essential for animals to maintain their life such as growth, metabolism, responsiveness, and movement. It is therefore important to understand how animal cells trigger hypoxia response and adapt to hypoxia thereafter. Both mammalian vascular and insect tracheal branches are induced to enhance the oxygen delivery. However, the study of hypoxia response in the nervous system remains limited. In this study, we assess the morphology of *Drosophila* neuromuscular junctions (NMJs), a model system to study development and function of synapses, in two hypoxia conditions, one with raising wild-type larvae in hypoxia, and the other in the *trachealess* (*trh*) mutant in which the trachea is defective, causing insufficient oxygen supply. Interestingly, glia, normally wrapping the axons of NMJs, invade into synapse and trigger Wg signals to reconstitute the synaptic structure under hypoxia. This synaptic remodeling maintains the synaptic transmission of synapse, which associate the locomotor behavior of larvae.

## Introduction

Animals need oxygen and food, not only to sustain life, but also for motility. In vertebrates, oxygen and nutrients are delivered through the vascular systems to organs and tissues throughout the body. To maintain proper nutrient and oxygen supply, and thus physiological functions, the vascular system is also highly coordinated with the nervous system during development. Indeed, the vascular and nervous systems resemble each other in terms of their anatomical structures and developmental processes [1, 2]. In the brain, nerves and vessels, form close associations and are in physical contact through the third player astrocytes to form neurovascular units (NVU) [3]. Such organization is essential for controlling oxygen and glucose delivery through the blood vessels by neuronal activity, and this regulatory process is mediated through the coupled astrocytes [4]. However, some invertebrates lack the complex vascular systems [5]. In nematodes, oxygen is supplied simply by ambient diffusion to inner cells [6]. Insects such as *Drosophila* have evolved a prototype of the tracheal system to deliver oxygen and a primitive vascular system, the dorsal vessel, to facilitate nutrient delivery [7]. However, the physical association of nerves, trachea, and glial processes has also been demonstrated at the NMJs of adult *Drosophila* flight muscles [8].

Animals respond to changing oxygen levels by altering their oxygen delivery system. Insufficient oxygen levels (hypoxia) activate a broad range of genes to re-establish body homeostasis. One crucial regulator of these hypoxia-responsive genes is the sequence-specific DNA-binding transcription factor hypoxia inducible factor 1 (HIF-1) [9]. HIF-1 consists of α and β subunits that form heterodimers [10]. Whereas HIF-1β is expressed constitutively, HIF-1α protein levels are modulated by oxygen levels [11]. Under normal oxygen conditions (normoxia), oxygen-dependent prolyl hydroxylases (PHDs) catalyze hydroxylation of a conserved prolyl residue in the central oxygen-dependent degradation (ODD) domain of HIF-1α [12-14]. Hydroxylation of HIF-1α promotes interaction with Von Hippel Lindau (VHL), which is the substrate recognition subunit of the cullin2-based E3 ubiquitin ligase, leading to HIF-1α ubiquitination and proteasomal degradation [15]. Under hypoxia, prolyl hydroxylation does not occur, HIF-1α proteins are stabilized and are translocated from the cytoplasm to the nucleus where they form heterodimers with HIF-1β to activate transcription of target genes [16, 17]. One major class of target genes encoding the Fibroblast Growth Factor (FGF) is involved in inducing angiogenesis in mammals. In *Drosophila,* the FGF member encoded by *Branchless* (*Bnl*) induces tracheal branching [18]. When oxygen levels are reduced, oxygen-starved cells express Bnl as a chemo-attractant to guide the growth tracheal terminal branches toward them [19].

In addition to adaptations of the respiratory system, the nervous system also responds to hypoxia. Oxygen levels modulate the survival, proliferation, and differentiation of radial glial cells (RGCs) in the human cerebral cortex. Interestingly, physiological hypoxia (3% O_2_) induces neurogenesis and differentiation of RGCs into glutamatergic neurons [20]. Hypoxia induces neurite outgrowth in PC12 cells through activation of A2A receptor [21]. Brief exposure to anoxia and hypoglycemia caused axonal remodeling in hippocampal neurons, including presynaptic protrusion of filopodia and formation of multi-innervated spines [22]. Under hypoxia or upon depletion of PHD2, upregulation of the actin cross-linker Filamin A (FLNA) induces generation of more immature spines [23]. Astrocytes have been shown to play a crucial role in ischemic tolerance via the activation of P2X7 receptors, which trigger upregulation of HIF-1α [24].

Neuronal PAS (NPAS) proteins containing a DNA-binding Per-Arnt-Sim domain function in vascular and nervous system development. In mice, NPAS1 is responsible for cortical interneuron generation [25], whereas NPAS3 is required for adult neurogenesis [26]. NPAS1 and NPAS3 also play key roles in lung development [27, 28]. The homolog of NPAS1/3 in *Drosophila*, Trachealess (Trh), has been well studied for its involvement in formation of the respiratory tracheal system. Trh is a master regulator of tracheal cell fates, activating gene expression to induce tracheal development [29, 30]. However, the role of Trh in the development of other tissues, particularly the nervous system, is unknown. In this study, we found altered synaptic bouton morphology at the NMJs of *trh*^*1*^*/trh*^*2*^ mutants. By performing *trh-RNAi* knockdown and *UAS-trh* transgene rescue experiments, we show that *trh* is required in tracheal cells for normal bouton formation. Defective tracheal branching in the *trh*^*1*^*/trh*^*2*^ mutant mimics the effect of hypoxic conditions during larval development, and supplying higher than normal oxygen levels restored normal bouton morphology. We further show that glial cells respond to hypoxia by elevating Wnt/Wg expression to mediate synaptic bouton remodeling through HIF1-α/Sima in *Drosophila*. Finally, we reveal that this synaptic remodeling maintains normal synaptic transmission and it is required for normal locomotion in larvae.

## Results

### *trh* modulates synaptic bouton formation non-cell autonomously

To understand the possible role of Trh in synapse formation, we examined NMJ morphology in the *trh* mutants. Since both *trh*^*1*^ *and trh*^*2*^ loss-of-function alleles are homozygous lethal [31-33], we examined the trans-heterozygous *trh*^*1*^*/trh*^*2*^ mutant that survive to adult stages and compared it to wild-type (*w*^*1118*^) and heterozygous *trh*^*1*^*/+* controls. Synaptic boutons of *w*^*1118*^ and *trh*^*1*^*/+* NMJs were evenly spaced along the axonal terminals, displaying the typical “beads-on-a-string” pattern (Fig 1A, upper and middle panels, enlarged images at right). Strikingly, the *trh*^*1*^*/trh*^*2*^ mutant larvae exhibited aberrant NMJ morphology in that their synaptic boutons were small and formed clusters without discernable connections, particularly at the terminals (Fig 1A, bottom panel); a phenotype described as “bunch boutons” [34]. This bunch bouton phenotype in the *trh*^*1*^*/trh*^*2*^ mutant was detected at a high frequency; 18% of total boutons were bunched compared to 3% for *trh*^*1*^*/+* and 0% for *w*^*1118*^ (Fig 1B). Of the 12 *trh*^*1*^*/trh*^*2*^ NMJs we examined, 11 possessed at least one bunch, with an average of 5 bunches per NMJ. Each bunch consisted of 3 to 10 small boutons (mean = 4.3). The *trh*^*1*^*/+* larvae exhibited a much weaker phenotype; only 4 of 10 examined larvae had 1 or 2 bunches, with an average of 5 small boutons per bunch. We did not observe a bunch bouton phenotype in any of the nine *w*^*1118*^ NMJs we assayed (Fig 1B). Although the percentage of bunch boutons in the *trh*^*1*^*/trh*^*2*^ mutant was greatly increased, total bouton number was only slightly higher than that observed in controls, suggesting that bunch boutons form at the expense of normal ones (Fig 1B). We also assessed the percentage of satellite boutons that are also small ones stemmed from normal-size boutons, and are often observed in wild-type NMJs. We observed some satellite boutons in the *w*^*1118*^ control and the *trh*^*1*^*/trh*^*2*^ mutant, and found no significant differences between them (Fig S1A). Given the small size and clustering of synaptic boutons in the *trh*^*1*^*/trh*^*2*^ mutant, we examined whether these bunch boutons express synaptic proteins normally. We found that the synaptic vesicle protein Synapsin (Syn in Fig 1A) was normally distributed relative to control, but the active zone protein Bruchpilot (Brp) was expressed at higher levels in bunch boutons (Fig S1B). The postsynaptic glutamate receptor, as revealed by GluRIIA (Fig S1C) and GluRIII (Fig S1B) signal, as well as dPAK (Fig S1C) were also localized in bunch boutons, which were surrounded by the subsynaptic reticulum protein Dlg (Fig S1D). Thus, although the Brp signal intensity in the *trh*^*1*^*/trh*^*2*^ mutant was stronger than in the wild-type, the composition of synaptic proteins in bunch boutons was largely similar to that of normal-sized boutons.

**Fig 1.**
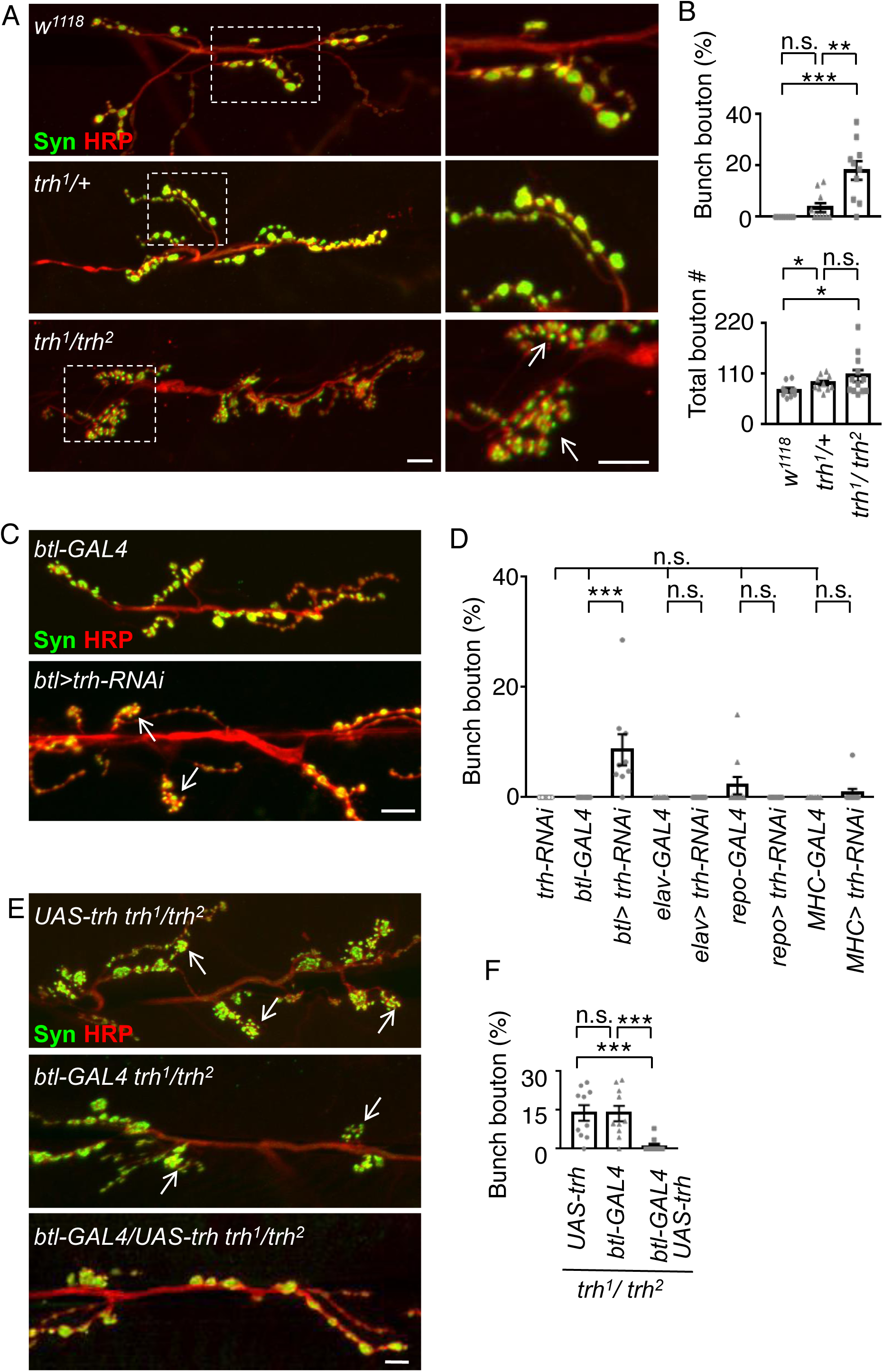
Bunch bouton formation at *trh* NMJs. (A, C, E) Immunostaining images show NMJs of muscle 6/7 for Syn (green) and HRP (red) in *w*^*1118*^, *trh*^*1*^*/+*, and *trh*^*1*^*/trh*^*2*^ (A), *btl-GAL4* and *btl*-*GAL4/trh-RNAi* (C), and *UAS-trh trh*^*1*^*/trh*^*2*^, *btl-GAL4 trh*^*1*^*/trh*^*2*^, *btl-GAL4/UAS-trh trh*^*1*^*/trh*^*2*^ (E). Dashed squares in (A) are enlarged in right panels. White arrows indicate clusters of bunch boutons. Scale bars are 10 μm. (B, D, F) Bar graphs with data dots within each bar show percentages (mean ± standard error of mean, SEM) of bunch boutons to total boutons. (B) *w*^*1118*^, 0 ± 0%, n = 10; *trh*^*1*^*/+*, 3.64 ± 1.74%, n = 10; *trh*^*1*^*/trh*^*2*^, 18.04 ± 3.65%, n = 10. (D) *trh-RNAi,* 0 ± 0%, n = 10; *btl-GAL4*, 0 ± 0%, n = 10; *btl>trh-RNAi*, 8.64 ± 2.81%, n = 9; *elav-GAL4*, 0 ± 0%, n = 10; *elav>trh-RNAi*, 0 ± 0%, n = 10; *repo-GAL4*, 2.14 ± 1.56%, n = 10; *repo*>*trh-RNAi*, 0 ± 0%, n = 10; *MHC-GAL4*, 0 ± 0%, n = 10; *MHC>trh-RNAi*, 0.77 ± 0.77%, n = 10. (F) *UAS-trh trh*^*1*^*/trh*^*2*^, 13.80 ± 3.00%, n = 10; *btl-GAL4 trh*^*1*^*/trh*^*2*^, 13.56 ± 2.95%, n = 10; *btl-GAL4/UAS-trh trh*^*1*^*/trh*^*2*^, 0.88 ± 0.64%, n = 8. (B, bottom panel) Total bouton numbers in *w*^*1118*^ (74.10 ± 4.72, n = 10), *trh*^*1*^*/+* (89.60 ± 5.16, n = 10), and *trh*^*1*^*/trh*^*2*^ (95.00 ± 11.21, n = 9). Statistical significance by Mann-Whitney test is shown as n.s., no significance; *, p < 0.5; **, p < 0.01; ***, p < 0.001.

As *trh* is expressed in both tracheal and nervous systems during embryonic stages [35], altered bouton morphology in the *trh*^*1*^*/trh2* mutant could be due to a lack of *trh* in neurons, tracheal cells or other cells/tissues. Therefore, we performed *trh-RNAi* knockdown by using tissue-specific GAL4 drivers for trachea (*btl-GAL4*), neurons (*elav-GAL4*), glia (*repo-GAL4*), and muscles (*MHC-GAL4*). We observed a dramatic increase in bunch boutons upon tracheal *trh* knockdown using *btl-GAL4* (Fig 1C and 1D). In contrast, *trh-RNAi* alone or using *elav-GAL4, repo-GAL4* or *MHC-GAL4* failed to replicate the bunch bouton phenotype (Fig 1D).

To further confirm the necessity of tracheal *trh* for normal bouton formation, we performed a rescue experiment of the *trh*^*1*^*/trh*^*2*^ phenotype. When we expressed a *UAS-trh* transgene in the trachea of the *trh*^*1*^*/trh*^*2*^ mutant upon using the tracheal *btl-GAL4* driver, the bunch bouton phenotype was suppressed (Fig 1E and 1F). Controls bearing only the *btl-GAL4* driver or the *UAS-trh* transgene still contained the comparably high numbers of bunch boutons observed in *trh*^*1*^*/trh*^*2*^ (Fig 1B) or upon tracheal *trh-RNAi* knockdown (Fig 1D). These results indicate that *trh* is required in the trachea for normal bouton formation.

### Hypoxia induces bunch bouton formation

Apart from specifying the tracheal cell fate, Trh is also involved in the branching of tubular structures during post-embryonic stages [30]. Therefore, we examined the tracheal phenotypes in the *trh*^*1*^*/trh*^*2*^ larvae and observed an increase of the number of terminal branches in the dorsal branch of the third segment (Fig S2A and S2B). Furthermore, we identified morphological defects such as tracheal breaks and tangles, suggesting structural defects in the *trh*^*1*^*/trh*^*2*^ larvae (arrows in Fig S2A). Tracheal branching activity is enhanced under hypoxia [18]. Thus, the increased number of terminal branches in *trh*^*1*^*/trh*^*2*^ could be a compensatory mechanism for defective trachea formation.

To understand whether *trh*^*1*^*/trh*^*2*^ mutant cells are under hypoxia, we used the hypoxia biosensor GFP-ODD, in which the GFP is fused to the oxygen-dependent degradation (ODD) domain of Sima, under the control of the *ubiquitin-69E* (*ubi*) promoter [36]. We first confirmed that GFP-ODD signal was low under normoxia (21% O_2_) and enhanced under hypoxia (5% O_2_, Fig 2A) in wild-type late-stage embryos when tracheal tubules are already formed and functioning [36]. Indeed, enhanced GFP signal was ubiquitous under hypoxia in wild-type embryos, with some pronounced focal GFP signals (Fig 2A, upper row, and Fig S2C). The signal of mRFP-nls, also under the control of *ubi* promoter as an internal control, remained constant under hypoxia (Fig 2A, bottom row, and Fig S2D) [36]. Quantification of the GFP/RFP ratio revealed a significant difference between normoxia and hypoxia conditions (Fig 2B). We then examined whether oxygen supply is deficient in the *trh*^*1*^*/trh*^*2*^ mutant by measuring the GFP-ODD signals. We detected higher GFP-ODD signal under hypoxia in the mutant compared to *w*^*1118*^ control (Fig 2A and Fig S2C). The heterozygous *trh*^*1*^*/+* presented similar GFP-ODD signal to *w*^*1118*^. Quantification of the GFP/RFP ratio also demonstrated a significant increase in GFP-ODD signal in *trh*^*1*^*/trh*^*2*^ (Fig 2B), supporting that the *trh*^*1*^*/trh*^*2*^ mutant senses reduced oxygen levels.

**Fig 2.**
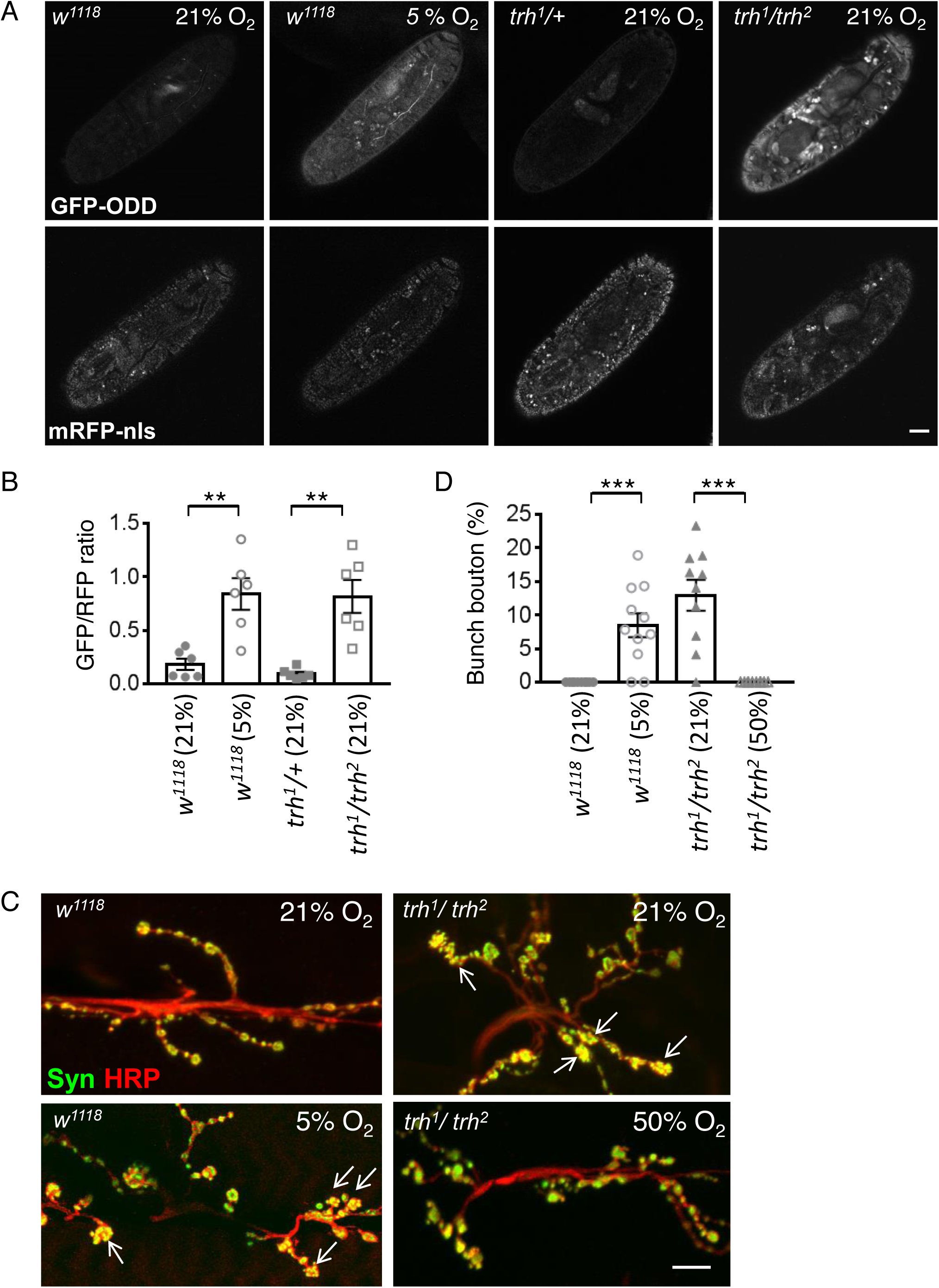
Induction of bunch boutons in *trh* mutants by hypoxia. (A) Images of late-stage embryos of *w*^*1118*^, *trh*^*1*^*/+*, and *trh*^*1*^*/trh*^*2*^ carrying the *GFP-ODD* reporter (top row) and *mRFP-nls* control (bottom row) under normoxia (21% O_2_) or hypoxia (5% O_2_). Scale bar represents 40 μm. (B) Bar graphs show mean ± SEM of the GFP/RFP immunofluorescence intensity ratio (*w*^*1118*^ at 21% O_2_, 0.18 ± 0.05, n = 6; *w*^*1118*^ at 5% O_2_, 0.84 ± 0.15, n = 6; *trh*^*1*^*/+* at 21% O_2_, 0.09 ± 0.02, n = 6; *trh*^*1*^*/trh*^*2*^ at 21% O_2_, 0.82 ± 0.15, n = 6). Statistical significance by Mann-Whitney test is shown as n.s., no significance; **, p < 0.01; ***, p < 0.001. (C) Images of NMJs from muscle 6/7 immunostained for Syn (green) and HRP (red) from *w*^*1118*^ at 21% or 5% O_2_ (left panels) and *trh*^*1*^*/trh*^*2*^ (right panels) at 21% or 50% O_2_. White arrows indicate clusters of bunch boutons. Scale bar represents 10 μm. (D) Bar graph shows percentages (mean ± SEM) of bunch boutons to total boutons (*w*^*1118*^ at 21% O_2_, 0 ± 0%, n = 10; *w*^*1118*^ at 5% O_2_, 8.47 ± 1.78%, n = 10; *trh*^*1*^*/trh*^*2*^ at 21% O_2_, 12.91 ± 2.30%, n = 10; *trh*^*1*^*/trh*^*2*^ at 50% O_2_, 0 ± 0%, n = 10). Statistical significance by Mann-Whitney test is shown as ***, p < 0.001.

Thus, formation of bunch boutons in the *trh*^*1*^*/trh*^*2*^ mutant could be caused by hypoxia. To test this hypothesis, we reared wild-type larvae under hypoxia (5%) and assessed synaptic bouton morphology at the third instar stage. Consistently, we observed small clustered boutons, mirroring the bunch bouton phenotype, at NMJs (Fig 2C and 2D). Furthermore, when we subjected the *trh*^*1*^*/trh*^*2*^ mutant to a high oxygen level (50%), the bunch bouton phenotype was completely suppressed, with *trh*^*1*^*/trh*^*2*^ NMJs exhibiting normal bouton morphology (Fig 2C and 2D). These results suggest that hypoxia due to the defective tracheal system in the *trh*^*1*^*/trh*^*2*^ mutant induces the bunch bouton phenotype, and that this phenotype can be suppressed by extra oxygen supply.

### **Glial HIF-1**α**/Sima mediates bunch bouton formation**

HIF-1α/Sima mediates the response to low oxygen supply [37, 38]. Protein levels of Sima are increased in wild-type *Drosophila* embryos subjected to hypoxia [39], leading to transcriptional activation of downstream target genes and the induction of tracheal branching [18]. We overexpressed Sima in tracheal cells, neurons, glia, or muscle cells by tissue-specific drivers to investigate which types of cells may play a role in modulating synaptic bouton formation. Overexpressing Sima in trachea caused embryonic lethality, preventing us from observing NMJ phenotypes. Larvae in which Sima was overexpressed in muscles, neurons, and glia could survive to the third instar stage and we detected a substantial number of bunch boutons upon glial Sima overexpression (Fig 3A and 3B). This result shows that the hypoxia-responding factor Sima is capable of inducing bunch bouton formation when overexpressed in glia.

**Fig 3.**
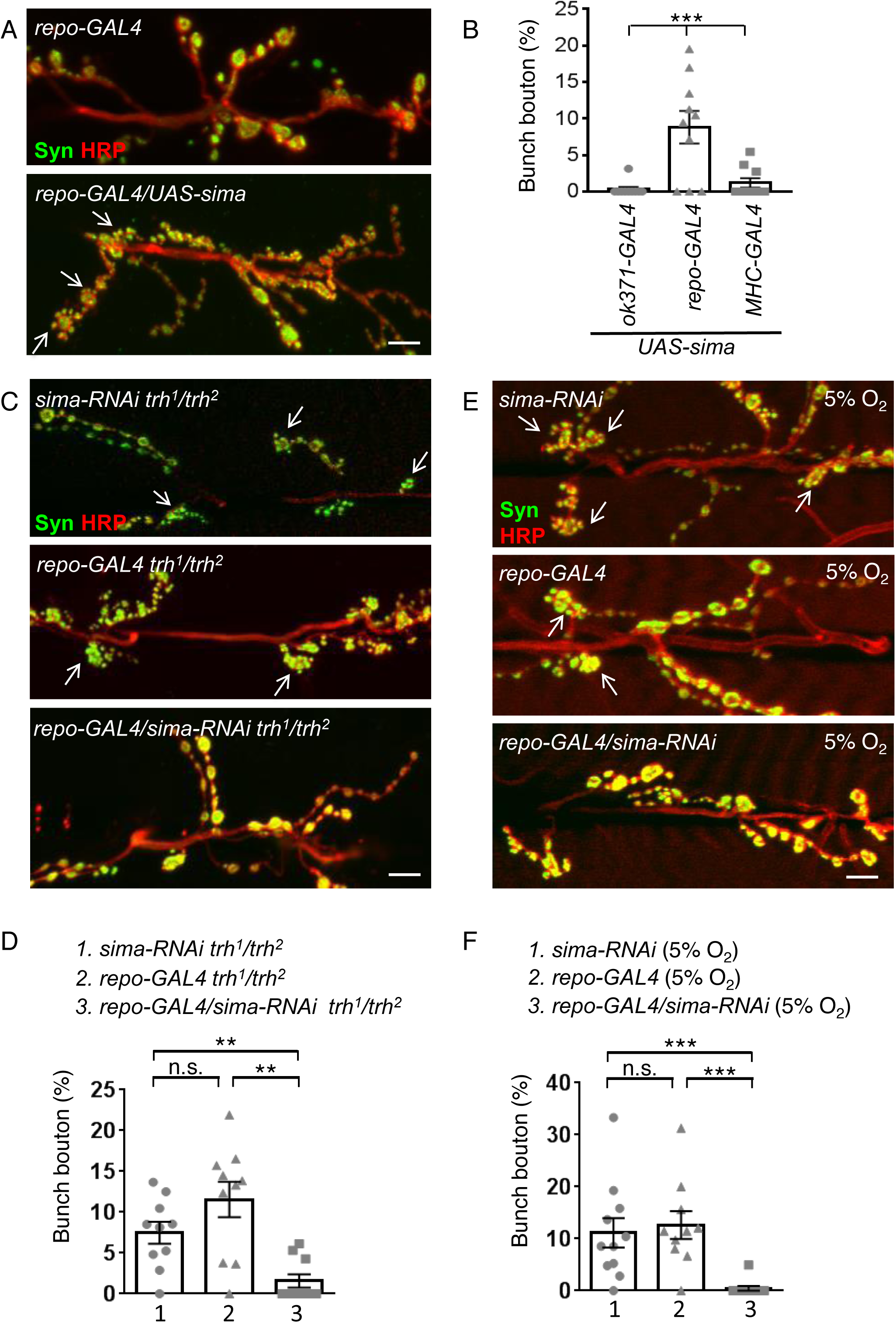
Bunch bouton induction by glial Sima at *trh* NMJs. (A, C, E) Images showing NMJs from muscle 6/7 immunostained for Syn (green) and HRP (red) in *repo-GAL4,* and *repo-GAL4/UAS-sima* (A), *sima-RNAi trh*^*1*^*/trh*^*2*^, *repo-GAL4 trh*^*1*^*/trh*^*2*^, and *repo-GAL4/sima-RNAi trh*^*1*^*/trh*^*2*^ (C), *sima-RNAi, repo-GAL4*, and *repo*-*GAL4*/*sima-RNAi* under hypoxia at 5% O_2_ (E). White arrows indicate bunch boutons. Scale bars represent 10 μm. (B, D, F) Bar graphs show percentages (mean ± SEM) of bunch boutons (B) *ok371-GAL4*/*UAS-sima*, 0.31 ± 0.31%, n = 10; *repo-GAL4*/*UAS-sima*, 8.81 ± 2.22%, n = 10; *MHC-GAL4*/*UAS-sima*, 1.19 ± 0.64%, n = 10. (D) *sima-RNAi trh*^*1*^*/trh*^*2*^, 7.48 ± 1.35%, n = 10; *repo-GAL4 trh*^*1*^*/trh*^*2*^, 11.56 ± 2.17%, n= 10; *repo-GAL4/sima-RNAi trh*^*1*^*/trh*^*2*^, 1.58 ± 0.81%, n = 10. (F) *sima-RNAi* at 5% O_2_,11.14 ± 2.8%, n = 11; *repo-GAL4* at 5% O_2_, 12.65 ± 2.66%, n = 10; *repo*-*GAL4*/*sima-RNAi* at 5% O_2_, 0.45 ± 0.45%, n = 11. Statistical significance by one-way ANOVA with Tukey’s Multiple Comparison post test is shown as ***, p < 0.001 (B) or Mann-Whitney test as n.s., no significance; **, p < 0.01; ***, p < 0.001 (D, F).

If glial Sima is the factor responsible for responding to hypoxia in the *trh*^*1*^*/trh*^*2*^ mutant, reducing the Sima level in glia would suppress bunch bouton formation. Accordingly, we expressed the *sima-RNAi* transgene, which could effectively deplete *sima* expression (Fig S3A), using *repo-GAL4* in the *trh*^*1*^*/trh*^*2*^ mutant. As our prediction, the bunch bouton phenotype was almost completely suppressed upon glial *sima* knockdown (Fig 3C and 3D). In controls, *trh*^*1*^*/trh*^*2*^ mutants carrying only the *UAS-sima-RNAi* transgene or the *repo-GAL4* driver still exhibited large numbers of bunch boutons (Fig 3C and 3D). We also tested whether low oxygen level-induced bunch bouton formation is mediated through Sima in glia. Bunch bouton phenotypes were detected in controls carrying either *repo-GAL4* or *UAS-sima-RNAi* when raised in 5% O_2_ (Fig 3E). However, almost no bunch boutons were detected in larvae carrying both *repo-GAL4* and *UAS-sima-RNAi* when raised in the same low-oxygen environment (Fig 3E and 3F). We also examined Sima protein levels in larvae, and found ubiquitous increases (including in Repo-positive glia) in the *trh*^*1*^*/trh*^*2*^ mutant or for the control under the 5% O_2_ condition (Fig S3B). Thus, glial Sima mediates the hypoxia response in the *trh*^*1*^*/trh*^*2*^ mutant and in the low O_2_ condition to modulate synaptic bouton formation.

### Glial Wg remodels bouton morphology

Next, we explored possible signals transduced from glia to neurons in response to hypoxia. The glia-secreted Wingless (Wg) signaling molecule regulates synaptic growth at *Drosophila* NMJs [40, 41]. Therefore, we examined whether Wg can be induced under hypoxia in the *trh*^*1*^*/trh*^*2*^ mutant. Wg signal was enriched around the synaptic boutons of wild-type NMJs (Fig 4A), similar to previous reported staining patterns [40, 42]. Although the pattern of Wg signal at *trh*^*1*^*/+* NMJs was similar to that of *w*^*1118*^, we detected much higher levels of Wg at *trh*^*1*^*/trh*^*2*^ NMJs (Fig 4A). Quantification of Wg immunofluorescence intensities normalized to co-stained HRP revealed a ∼3-fold increase relative to *w*^*1118*^ and *trh*^*1*^*/+*, respectively (Fig 4B). We then examined whether Sima is required for the enhanced Wg expression in the *trh*^*1*^*/trh*^*2*^ mutant. In a *trh*^*1*^*/trh*^*2*^ mutant carrying *repo-GAL4*, Wg levels were also increased 3.0-fold relative to the *repo-GAL4* control (Fig 4C, 4D). When we reduced *sima* levels in the *trh*^*1*^*/trh*^*2*^ mutant by *repo-GAL4*-driven *UAS-sima-RNAi*, Wg signal was suppressed to a level equivalent to that in the *repo-GAL4* control (Fig 4C and 4D). Interestingly, *sima-RNAi* knockdown in glia of the *repo-GAL4* control had no effect on the Wg level (Fig 4C and 4D), suggesting that Sima is induced to upregulate Wg expression in the *trh*^*1*^*/trh*^*2*^ mutant but has no role in basal Wg expression in the wild-type. Taken together, we suggest that glial Sima is required for Wg upregulation at the NMJs of the *trh*^*1*^*/trh*^*2*^ mutant.

**Fig 4.**
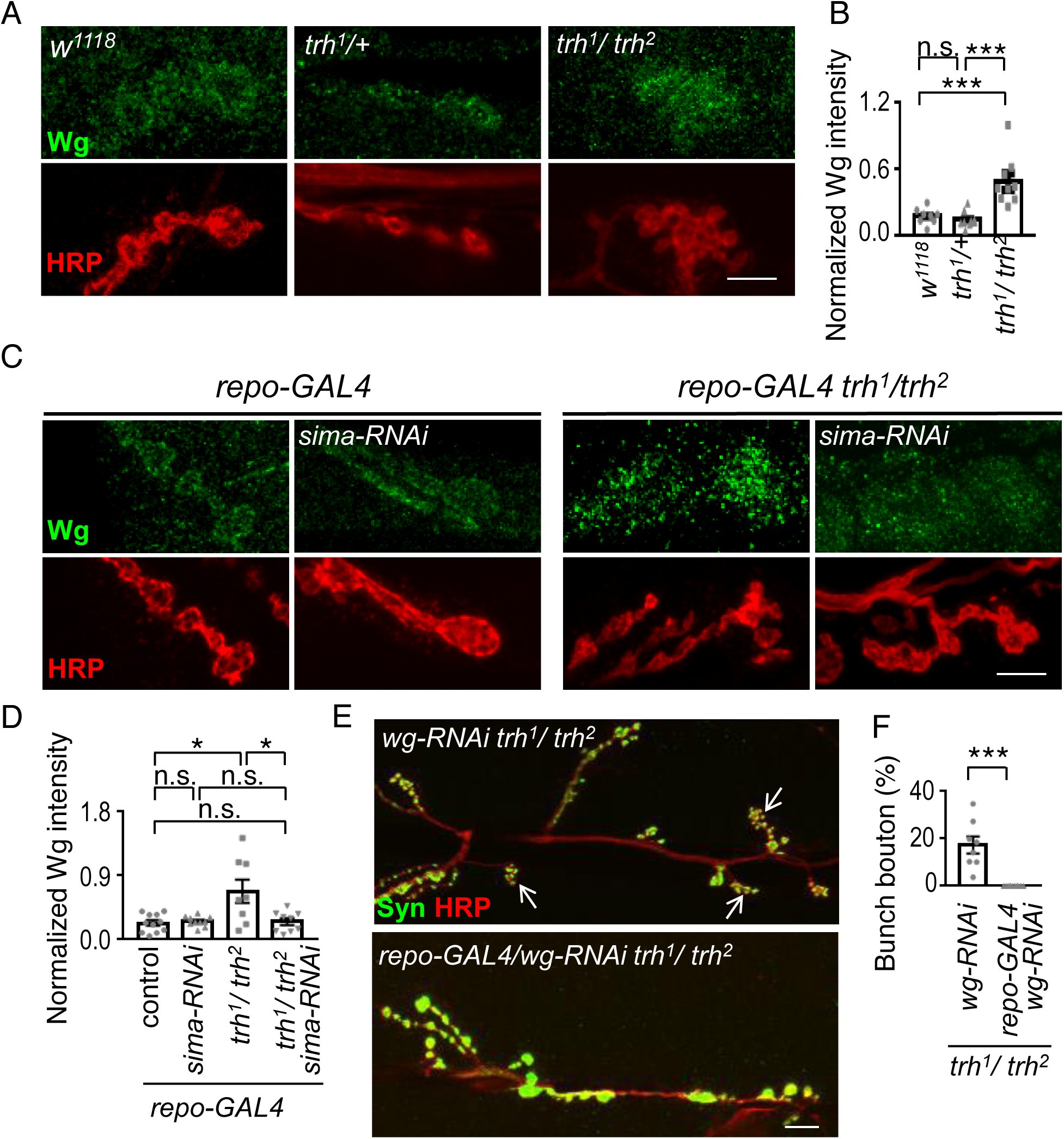
Wg induction from glia modulates synaptogenesis at *trh* NMJs. (A, C, E) Images showing NMJs of muscle 6/7 immunostained for Wg (green) and HRP (red) in *w*^*1118*^, *trh*^*1*^*/+*, and *trh*^*1*^*/trh*^*2*^ (A); *repo-GAL4, repo-GAL4*/*sima-RNAi, repo-GAL4 trh*^*1*^*/trh*^*2*^, and *repo-GAL4/sima-RNAi trh*^*1*^*/trh*^*2*^ (C); and for Syn (green) and HRP (red) in *wg-RNAi trh*^*1*^*/trh*^*2*^ and *repo-GAL4/wg-RNAi trh*^*1*^*/trh*^*2*^ (E). White arrows indicate clusters of bunch boutons. Scale bars represent 10 μm. (B, D, F) Bar graphs show averages (mean ± SEM) of normalized Wg to HRP intensities. *w*^*1118*^, 0.18 ± 0.03, n = 8; *trh*^*1*^*/+*, 0.15 ± 0.02, n = 9; *trh*^*1*^*/trh*^*2*^, 0.49 ± 0.07, n = 9 (B), *repo-GAL4*, 0.23 ± 0.04, n = 11; *repo-GAL4*/*sima-RNAi*, 0.25 ± 0.02, n = 10; *repo-GAL4 trh*^*1*^*/trh*^*2*^, 0.68 ± 0.16, n = 8; *repo-GAL4/sima-RNAi trh*^*1*^*/trh*^*2*^, 0.25 ± 0.05, n = 9 (D), and percentages (mean ± SEM) of bunch boutons in *wg-RNAi trh*^*1*^*/trh*^*2*^, 17.19 ± 3.52%, n = 8; *repo-GAL4/wg-RNAi trh*^*1*^*/trh*^*2*^, 0.68 ± 0.68%, n = 11 (F). Statistical significance by Mann-Whitney test is shown as n.s., no significance; *, p < 0.05; **, p < 0.01; ***, p < 0.001.

If glia-secreted Wg is responsible for bunch bouton induction in the *trh*^*1*^*/trh*^*2*^ mutant, then glia-specific knockdown of *wg* in that mutant should suppress bunch bouton formation. Indeed, the bunch bouton phenotype was almost undetectable in the *trh*^*1*^*/trh*^*2*^ mutant also bearing *repo-GAL4* and the *UAS-wg-RNAi* transgene (Fig 4E and 4F). In control, bunch boutons were still prominent in the *trh*^*1*^*/trh*^*2*^ mutant bearing only *UAS-wg-RNAi* (Fig 4E and 4F). Inactivation of Wg signaling has been shown to induce unbundled filaments and a reduction of the more stabilized loops upon immunostaining for the microtubule-binding protein Futsch [41, 43]. To show that elevated Wg signaling remodels presynaptic bouton structure in the *trh*^*1*^*/trh*^*2*^ mutant, we examined Futsch-labeled microtubules within synaptic boutons and found significantly more Futsch-positive loops within the boutons (Fig S4A and S4B), supporting the elevation of presynaptic Wg signaling. Taken together, these results suggest that Wg plays a prominent role in the *trh*^*1*^*/trh*^*2*^ mutant to transduce the hypoxia signal from glia to remodel presynaptic bouton structure.

Since glial processes can invade synaptic boutons to match the growth of NMJs [44], we were intrigued to assess whether glia the *trh*^*1*^*/trh*^*2*^ mutant exhibits morphological change. In a live preparation of NMJs, we found that glial processes invaded the area of synaptic boutons in the *trh*^*1*^*/trh*^*2*^ mutant, whereas glial processes were comparatively restrained in the control (Fig S4C). Quantification of fluorescent signals of glial process overlaying the synaptic bouton area revealed significantly greater area of overlap in the *trh*^*1*^*/trh*^*2*^ mutant relative to control (Fig S4D). This increased extent of glial processes in the synaptic area may facilitate signal transduction from glia to synaptic boutons for structural reorganization.

### Impaired crawling behavior in the *trh* mutant

Given the evident morphological changes at *trh*^*1*^*/trh*^*2*^ NMJs, we wondered if locomotion is affected in mutant larvae. We observed larvae crawling under free-movement conditions and found that wild-type control and *trh*^*1*^*/+* heterozygous larvae presented smooth crawling paths with an average speed of ∼0.5 mm/s (Fig 5A and 5B), whereas the *trh*^*1*^*/trh*^*2*^ larva had shorter paths and a slower speed of 0.14 mm/s. The head turning frequency in *trh*^*1*^*/trh*^*2*^ was comparable to both controls, not contributing to the slow moving (Fig 5C). Larval crawling is a rhythmic behavior involving a series of periodic strides (S1 Movie) [45-48]. We noticed uncoordinated crawling in *trh*^*1*^*/trh*^*2*^ larvae, with their posterior body segments failing to follow the rhythmic movement (S2 Movie). We recorded larval forward crawling and constructed kymographs to represent the stride cycle. In wild-type larvae, normal and consistent periodic strides were apparent with regular head and tail displacements (Fig 5D, left panel). Similar to the wild-type, head movements of *trh*^*1*^*/trh*^*2*^ larvae were smooth and periodic, albeit slower. However, tail movements of *trh*^*1*^*/trh*^*2*^ larvae were abrupt (Fig 5D, right panel). Approximately 70% of the strides of *trh*^*1*^*/trh*^*2*^ larvae were uncoordinated (Fig 5E), which might contribute to the slower crawling of the *trh*^*1*^*/trh*^*2*^ mutant.

**Fig 5.**
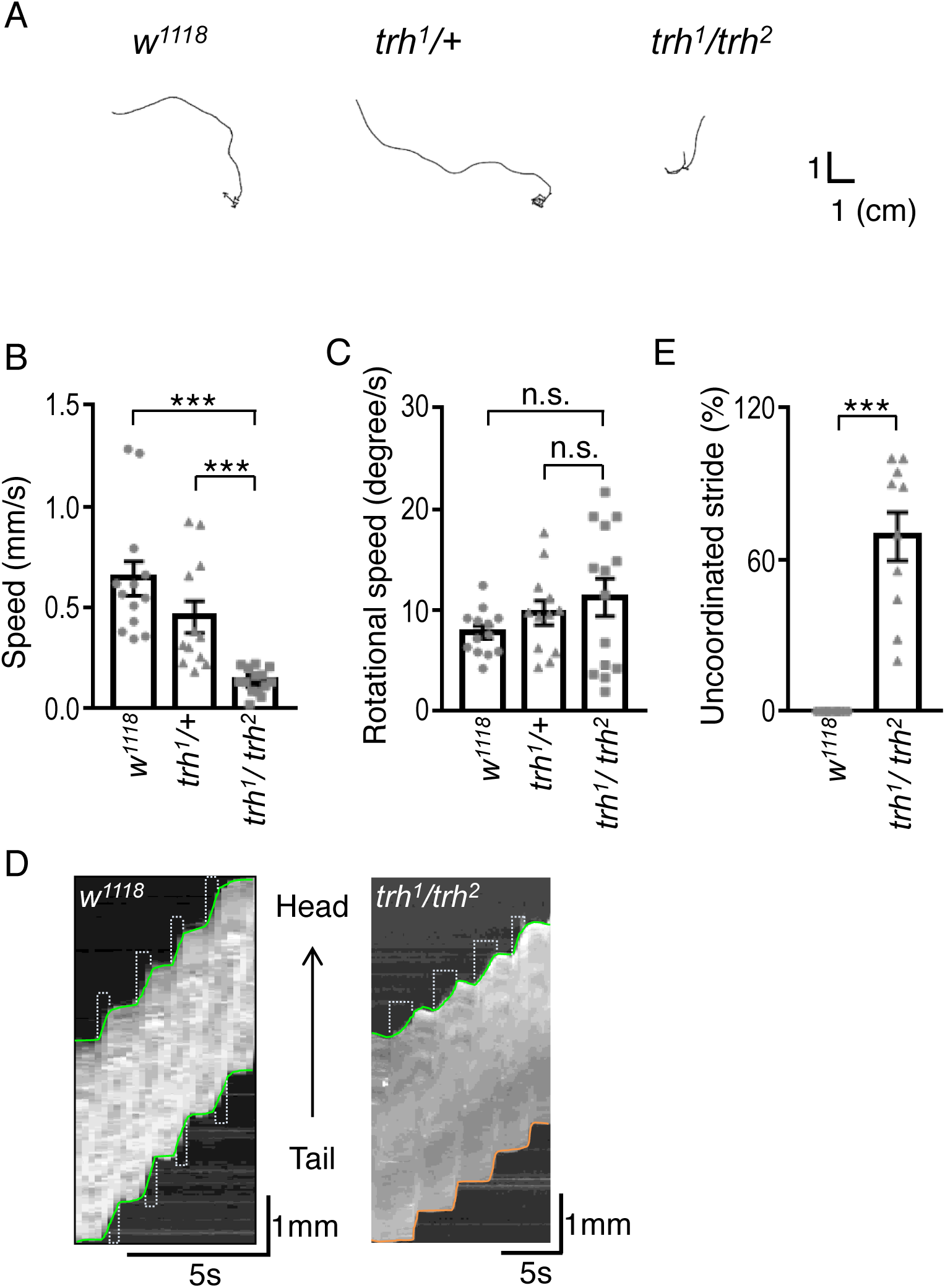
Impaired crawling behavior of the *trh* mutant. (A) Representative 5-minute crawling traces for *w*^*1118*^, *trh*^*1*^*/+*, and *trh*^*1*^*/trh*^*2*^. (B) Bar graphs show average speeds (mean ± SEM in mm/s) for *w*^*1118*^, 0.64 ± 0.09, n = 13; *trh*^*1*^*/+*, 0.45 ± 0.08, n = 12; *trh*^*1*^*/ trh*^*2*^, 0.14 ± 0.02, n = 14, and average rotational speeds (mean ± SEM in degree/s) for *w*^*1118*^, 7.80 ± 0.62, n = 13; *trh*^*1*^*/+*, 9.73 ± 1.19, n= 12; *trh*^*1*^*/trh*^*2*^, 11.26 ± 1.86, n = 14. Statistical significance by Mann-Whitney test is shown as n.s., no significance; ***, p < 0.001. (D) Kymographs show four larval forward crawling strides for *w*^*1118*^ and *trh*^*1*^*/trh*^*2*^. The green lines show smooth transitions between successive strides, whereas orange lines show abrupt transitions.(E) Bar graph shows uncoordinated stride percentages (mean ± SEM) in *w*^*1118*^ (0 ± 0%, n = 10) and *trh*^*1*^*/trh*^*2*^ (69.22 ± 9.57%, n = 10). Statistical significance by Mann-Whitney test is shown as ***, p < 0.001.

This uncoordinated stride cycle prompted us to examine the bouton morphology in anterior and posterior segments of *trh*^*1*^*/trh*^*2*^ larvae. Strikingly, we detected the bunch bouton phenotype in segments A2-A4 of individual *trh*^*1*^*/trh*^*2*^ mutant larvae, whereas bouton morphology in their respective posterior A5 and A6 segments was relatively normal (Fig 6A). In fact, we seldom observed the bunch bouton phenotype in posterior segments (Fig 6B). In wild-type larvae, all segments we examined had normal bouton morphology (Fig 6A and 6B). While bunch boutons appeared in A2-A4 segments, the total numbers of boutons were comparable in all segments between wild-type and *trh*^*1*^*/trh*^*2*^. We further examined whether Wg signal from glia has any effect on modifying bouton morphology in *trh*^*1*^*/trh*^*2*^ larvae. Glial *wg-RNAi* knockdown suppressed bunch bouton formation in the anterior A3 segment, but had no effect on the morphological phenotype of the posterior A6 segment (Fig 6E). Thus, induction of bunch boutons is specific to anterior A2-A4 segments.

**Fig 6.**
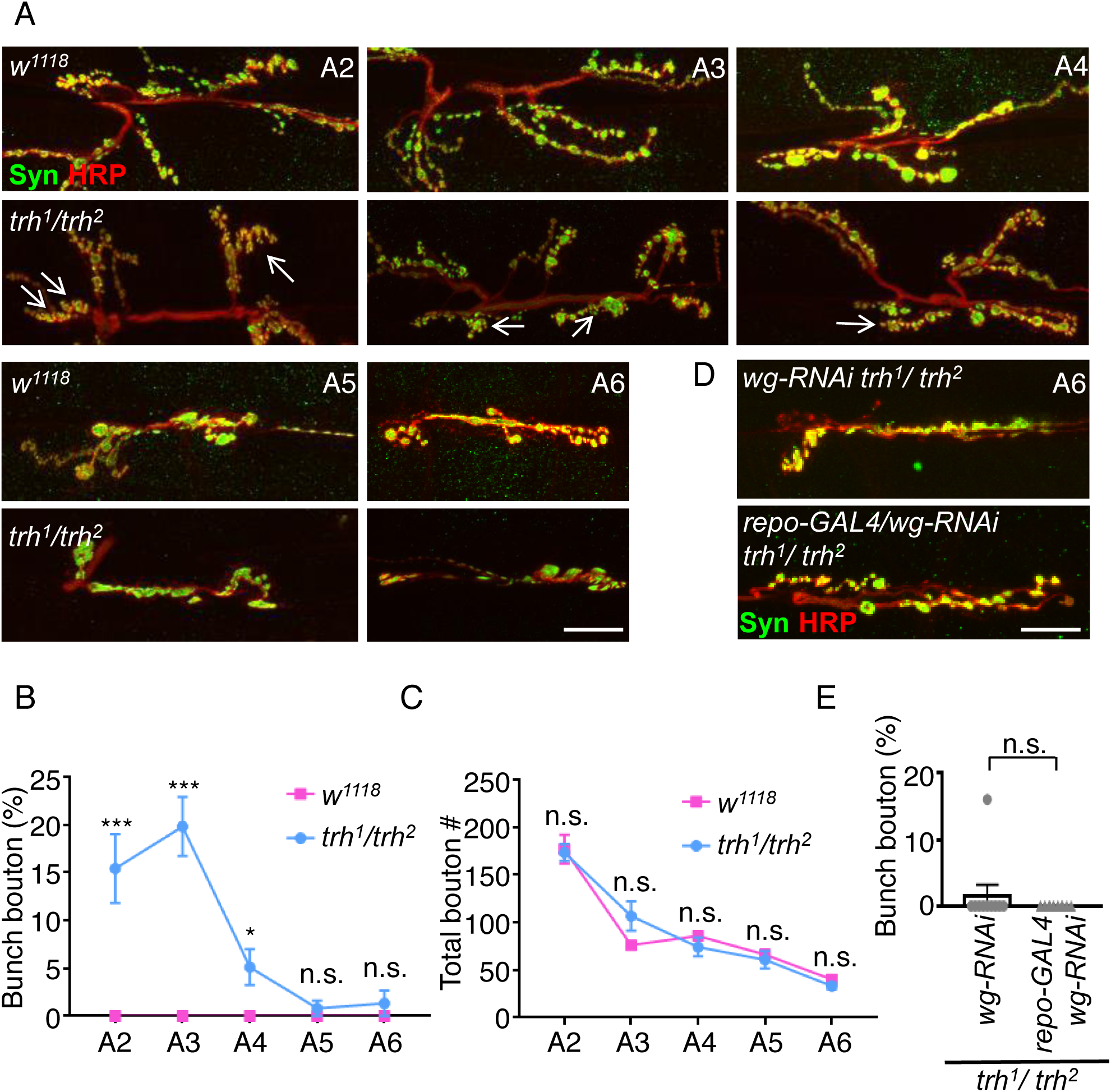
Selective bunch bouton formation at *trh* NMJs of anterior segments. (A) Images show NMJs of muscle 6/7 from abdominal segments A2-A6 immunostained for Syn (green) and HRP (red) in *w*^*1118*^ and *trh*^*1*^*/ trh*^*2*^. White arrows indicate clusters of bunch boutons. Scale bar represents 10 μm. (B, C) Line graphs show average percentages (mean ± SEM) of (B) bunch bouton (*w*^*1118*^: A2, 0 ± 0%, n = 10; A3, 0 ± 0%, n = 9; A4, 0 ± 0%, n = 10; A5, 0 ± 0%, n = 10; A6, 0 ± 0%, n = 9; *trh*^*1*^*/trh*^*2*^: A2, 15.40 ± 3.59%, n = 10; A3, 19.80 ± 3.09%, n = 10; A4, 5.11 ± 1.86%, n= 9; A5, 0.80 ± 0.80%, n = 10; A6, 1.33 ± 1.33%, n = 9) and total bouton numbers (*w*^*1118*^: A2, 176.60 ± 15.00, n = 10; A3, 76.11 ± 4.77, n = 9; A4, 85.80 ± 4.13, n = 10; A5, 66.10 ± 5.39, n = 10; A6, 40.00 ± 3.72, n = 9; *trh*^*1*^*/trh*^*2*^: A2, 173.20 ± 8.76, n = 10; A3, 106.60 ± 15.33, n = 10; A4, 74.11 ± 9.90, n = 9; A5, 60.70 ± 9.10, n = 10; A6, 33.11 ± 3.56, n = 9) for *w*^*1118*^ (magenta lines) and *trh*^*1*^*/ trh*^*2*^ (blue lines). Statistical significance by Mann-Whitney test is shown as n.s., no significance; *, p < 0.05; ***, p < 0.001. (D) Images showing NMJs of muscle 6/7 immunostained for Syn (green) and HRP (red) in *wg-RNAi trh*^*1*^*/trh*^*2*^ (top panel) and *repo-GAL4/wg-RNAi trh*^*1*^*/trh*^*2*^ (bottom panel) in A6. (D) Bar graph shows percentages (mean ± SEM) of bunch boutons (*wg-RNAi trh*^*1*^*/trh*^*2*^, 1.60 ± 1.60%, n = 10; *repo-GAL4/wg-RNAi trh*^*1*^*/trh*^*2*^, 0 ± 0%, n = 10). Statistical significance by Mann-Whitney test is shown as n.s., no significance.

Given the bunch bouton phenotype in *trh*^*1*^*/trh*^*2*^ larvae, we assessed basal synaptic transmission properties, firstly at muscle 6 of anterior A3 segments. Amplitude of evoked junctional potential (EJP), as well as the amplitude and frequency of miniature EJP (mEJP), were comparable among controls and *trh*^*1*^*/trh*^*2*^ larvae (**Fig 7A-7E**). The quantal content, calculated by dividing the EJP amplitude with that of mEJP, were also equivalent (**Fig 7F**). We then evaluated the synaptic transmission properties of muscle 6 for posterior A6 segments. Although the amplitude and frequency of mEJP and the EJP amplitude in *trh*^*1*^*/trh*^*2*^ larvae remained similar to controls, we did detect a significant reduction in the quantal content of mutant larvae (**Fig 7F**). Thus, the impaired synaptic activity of posterior segments of mutant larvae seems to be consistent with their defective stride cycle.

**Fig 7.**
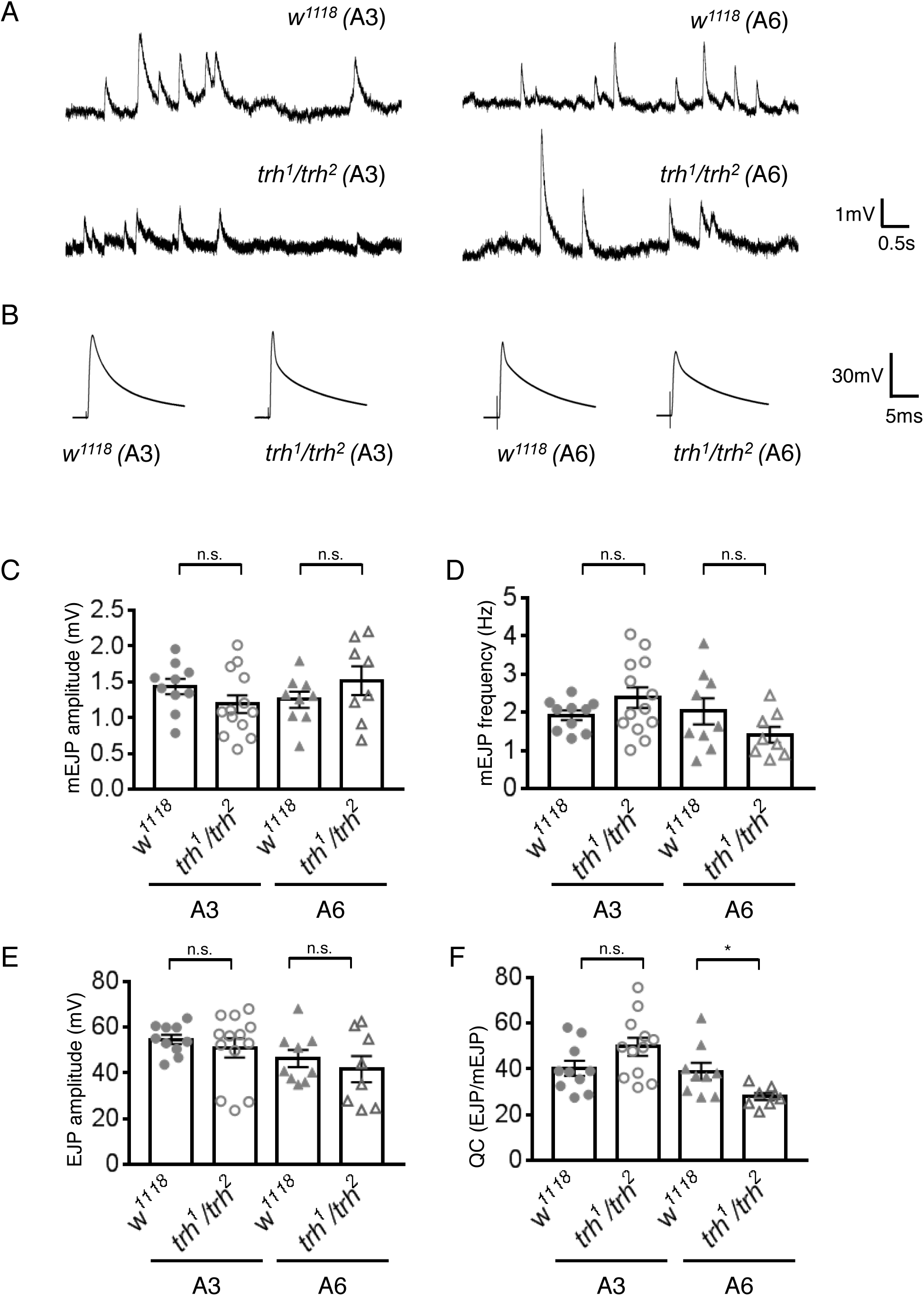
Synaptic transmission at *trh* NMJs. (A, B) Representative traces of mEJPs (A) and EJPs (B) recorded from NMJs of muscle 6 of segment A3 (left panels) and A6 (right panels) for *w*^*1118*^ and *trh*^*1*^*/trh*^*2*^. The scales for both mEJP and EJP traces are depicted at right. (C-F) Bar graphs show averages (mean ± SEM) for (C) mEJP amplitudes (*w*^*1118*^/A3, 1.43 ± 0.11, n = 10; *trh*^*1*^*/trh*^*2*^*/*A3, 1.19 ± 0.12, n = 13; *w*^*1118*^/A6, 1.25 ± 0.11, n = 9; *trh*^*1*^*/trh*^*2*^*/*A6, 1.51 ± 0.20, n = 8); (D) mEJP frequencies (*w*^*1118*^/A3, 1.92 ± 0.13, n = 10; *trh*^*1*^*/trh*^*2*^*/*A3, 2.38 ± 0.27, n = 13; *w*^*1118*^/A6, 2.02 ± 0.34, n = 9; *trh*^*1*^*/trh*^*2*^*/*A6, 1.41 ± 0.21, n = 8); (E) EJP amplitudes (*w*^*1118*^/A3, 54.61 ± 2.06, n = 10; *trh*^*1*^*/trh*^*2*^*/*A3, 50.89 ± 4.18, n = 13; *w*^*1118*^/A6, 46.24 ± 3.71, n = 9; *trh*^*1*^*/trh*^*2*^*/*A6, 41.64 ± 5.71, n = 8); and (F) quantal contents (QC) calculated as EJP/mEJP (*w*^*1118*^/A3, 40.23 ± 3.33, n = 10; *trh*^*1*^*/trh*^*2*^*/*A3, 49.63 ± 3.91, n = 13; *w*^*1118*^/A6, 38.90 ± 3.75, n = 9; *trh*^*1*^*/trh*^*2*^*/*A6, 27.88 ± 1.53, n = 8). Statistical significance by Mann-Whitney test is shown as n.s., no significance; *, p < 0.05; **, p < 0.01.

## Discussion

Here, we demonstrate that Trh, a member of the NPAS protein family, non-cell autonomously regulates synaptic bouton formation at NMJs through a hypoxia response from glia. We observed small-sized and clustered boutons, so-called bunch boutons, at the NMJs of *trh* mutant larvae or larvae reared at low oxygen levels. The abnormal bouton morphology at *trh* NMJs could be suppressed by reducing levels of the hypoxia response factor Sima in glia. We further showed that Sima enhanced the Wg signal from glia to cause bunch bouton formation. Although normal synaptic transmission was detected at NMJs located in anterior segments of larvae bearing bunch boutons, reduced synaptic transmission was found in posterior segments lacking bunch boutons of the *trh* mutant, suggesting that glia-induced bunch bouton formation might be a homeostatic response to restore normal synaptic transmission. Imbalanced synaptic functioning of mutant NMJs might contribute to the uncoordinated stride cycles detected in the *trh* mutant, slowing larval crawl speed. Thus, we provide a model for studying the glial responses that modulate synaptic remodeling during hypoxia.

### Glia play a critical role in response to hypoxia in the *trh* mutant

Animal cells adapt to hypoxia by triggering the expression of HIF-1α/Sima, the master transcriptional regulator of the hypoxia response [49]. We observed defective tracheal structure in the *trh* mutant (Fig S2A), which may result in hypoxic conditions inside the larval body. The increase in terminal branch number (Fig S2B) may be a response to oxygen supply deficiency [18]. Moreover, the increases in Sima protein levels and *ODD-GFP* reporter expression indicate reduced internal oxygen levels (Fig 2A, 2B and S3B). Finally, the bunch bouton phenotype in the *trh* mutant was recapitulated by rearing larvae under hypoxia, and it was suppressed by rearing larvae under hyperoxia. Taken together, these observations suggest that cells in the *trh* mutant sense low oxygen levels caused by the defective tracheal system and respond by elevating Sima protein levels. It is not clear how profound this effect is for other types of larval cells. Based on our ODD-GFP and Sima immunostaining patterns (Fig 2A, 2B and S3B), many types of cells are likely to be affected [50]. Our results also suggest that Trh has a late developmental role in tracheal morphogenesis, in addition to its well-characterized role in early tracheal cell fate specification [29, 30]. Unlike the mammalian homologs NPAS1 and NPAS3 that function in the nervous system, the *Drosophila* NPAS1/3 homolog Trh is more dedicated to tracheal development.

### Glia induce synaptic remodeling

We suggest that glia is the major cell type mediating bunch bouton formation in the *trh* mutant under hypoxia. HIF-1α/Sima was ubiquitously, including glia, increased in both the *trh*^*1*^*/trh*^*2*^ mutant and the control grown under hypoxia (Fig S3B). Importantly, by manipulating Sima expression in glia we could regulate bunch bouton formation (Fig 3). Elevated Sima levels induce tracheal sprouting in tracheal cells, as well as protrusions in non-tracheal cells [18]. Interestingly, we also observed increased overlap of glial processes with synaptic area in the *trh* mutant, indicative of a glial response (Fig S4C, S4D). Several types of cells in *Drosophila* have been shown to respond to hypoxia [39, 51]. For instance, under hypoxia, elevated Sima levels induce the expression of Breathless (Btl, the FGF receptor) in tracheal cells that branch out seeking cells that express Branchless (Bnl)/FGF, with this latter process also being partially dependent on Sima [18, 19]. In an alternative pathway, atypical soluble guanylyl cyclases that are less sensitive to nitric oxide than conventional soluble guanylyl cyclases can mediate graded and immediate hypoxia responses mainly in neurons [52, 53]. *Drosophila* glia have not been reported to sense and respond to hypoxia, but mammalian astrocytes in the central nervous system have been shown to be involved in these processes. In a middle cerebral artery occlusion mouse model, astrocyte activation was shown to play a crucial role in ischemic tolerance, which is mediated through P2X7 receptor-activated HIF-1α upregulation [24]. Under physiological hypoxia, reduced mitochondrial respiration leads to the release of intracellular calcium and exocytosis of ATP-containing vesicles, thereby signaling the brainstem to modulate animal breathing [54]. Our results reveal a role for *Drosophila* larval glia in sensing hypoxia via the conventional HIF-1α/Sima pathway, warranting further detailed study.

We also demonstrated that under hypoxia, glia modulate the formation of synaptic boutons (Fig 3). These results clearly place the glia-modulated morphology of synaptic boutons in the context of hypoxia responses. Several studies have suggested that glia play important roles in regulating synaptic morphology at developmental stages or in response to neural insult. Perisynaptic Schwann cells surround nerve terminals and express neurotransmitter receptors, modulating synaptic efficacy upon nerve stimulation at mouse NMJs [55]. After nerve injury, Schwann cells participate in synaptic homeostasis and remodeling during NMJ re-innervation [56]. Chronic hypoxia causes hypomyelination, leading to synaptic reduction in the mouse cortex, which could be rescued by genetically-induced or drug-enhanced hypermyelination [57]. In *Drosophila*, expansion of glial structures to synaptic boutons matches synaptic growth [44]. Glial invasion of the synaptic region is also suggested to clear presynaptic debris of unstable boutons during activity-dependent synaptic growth [58]. Our study further establishes that in response to hypoxia, Wg is a glial signal that modulates synaptic bouton formation. Two sources of Wg, from presynaptic motor neurons and from glia, are involved in synapse growth and remodeling [40]. Our results suggest that Sima upregulates the level of Wg secreted from glia to modulate synapse formation in the *trh* mutant or in control larvae grown under hypoxia. In hypoxic macrophages, HIF-1α mediates the induction of Wnt1, which is a mammalian homolog of Wg [59-61]. Although it is unclear whether Wg is a direct target of the hypoxia signal Sima in *Drosophila*, we show that the level of Wg is controlled by glial Sima (Fig 4C, 4D) and that it mediates Sima-induced bunch bouton formation at *trh* NMJs (Fig 4E, 4F). As a secreted morphogen, Wg functions in both pre-synaptic and post-synaptic sites [41]. At presynaptic terminals, the canonical Wg pathway induces microtubule loop formation to regulate synaptogenesis [43]. We also detected an increase in microtubule loops in the *trh* mutant (Fig S4A, S4B), consistent with a role for Wg in mediating the hypoxia response by modulating synapse formation. Postsynaptic Wg signaling leads to subsynaptic reticulum differentiation [41], which was not apparent in the *trh* mutant (Fig S1D), suggesting that Wg might be a component of the complex hypoxia response that induces synapse remodeling. Brief exposure to hypoxia induces immature spines and impaired synaptic function in hippocampal neurons [23]. The morphological change to bunch boutons at *trh* NMJs (Fig 1A and 1B) was not accompanied by altered synaptic transmission (Fig 7A-F), which may reflect a compensatory effect during long-term hypoxia.

The bunch bouton phenotype has also been described in *spastin* mutants [34, 62]. As an AAA ATPase, Spastin severs microtubules to facilitate transport to distal axon segments [63, 64]. Accordingly, the *spastin* mutant also exhibits a lack of microtubules at terminal boutons [62]. In contrast, the *trh* mutant presented an increase of microtubule loops (Fig S4A, S4B). Microtubule loops have been linked to synaptic bouton stabilization, and an excess of microtubule loops has been associated with increased synaptic bouton formation [65, 66]. The altered morphology of bunch boutons may be part of the structural changes necessary to maintain normal synaptic transmission under hypoxia. The *trh* and *spastin* mutants also exhibit differences in synaptic function, with loss of *spastin* function slightly enhancing spontaneous synaptic transmission release but reducing evoked synaptic transmission [62]. Thus, although the morphology of synaptic boutons at *trh* NMJs resembles that of *spastin* mutants, bunch boutons at *trh* NMJs retain synaptic functions, unlike the impaired synaptic transmission of *spastin* mutant boutons.

### Difference of *trh* NMJs in anterior and posterior segments

The size of NMJs in muscles 6/7 decreases from the anterior to posterior segments, which could represent a coupling with muscle growth [67, 68], thereby maintaining similar electrophysiological efficacy at anterior and posterior NMJs (Fig 7A-F). Interestingly, our findings show that synaptic responses in the *trh* mutant differ, with bunch boutons only appearing in anterior segments (Fig 6A and 6B). Furthermore, synaptic transmission at *trh* NMJs remained normal in anterior A3 segments but was impaired in posterior A6 segments (Fig 7A-F). These observations are consistent with bunch bouton formation being part of a homeostatic response to restore synaptic activity. Why synapses were not remodeled in posterior segments remains unclear. Motor neurons in the ventral nerve cord project much longer axons to muscles in posterior segments compared to anterior ones. It has been shown that axonal transport to posterior segments is more vulnerable to inefficient transport conditions. For example, mutation of long-chain Acyl-CoA synthetase impairs the balance between anterograde and retrograde transport, causing distally-biased axonal aggregations and affecting the growth and functioning of synapses [69]. In addition, larval forward locomotion, propelled by peristatic contraction, is controlled by different circuits targeting anterior and posterior segments. The GABAergic SEZ-LN1 neurons specifically control posterior A6 and A7 segmental muscle contraction by inhibiting A27h premotor neurons, which promotes longitudinal muscle contraction during larval forward crawling [70]. It is possible that glia-derived Wg signals from cell bodies located in the ventral nerve cord of the *trh* mutant may not be efficiently transported to posterior segments during hypoxia. This polar difference in synaptic activity and bouton morphology may contribute to the uncoordinated peristatic movements of *trh* larvae.

## Methods

### Fly stocks

All flies were reared at 25 °C. *w*^*1118*^ was used as wild-type control and to backcross with *trh*^*1*^ or *trh*^*2*^. Fly strains are as follows: *trh*^*2*^, *elav-GAL4, MHC-GAL4, repo-GAL4, ok371-GAL4, UAS-trh, UAS-sima*, and *UAS-sima-RNAi* from Bloomington Drosophila Stock Center (BDSC); *trh*^*1*^ and *UAS-wg-RNAi* from Kyoto Stock Center, and *UAS-trh-RNAi* from Vienna Drosophila Resource Center (VDRC). Also used were *btl-GAL4* [71], *GFP-ODD* [36], and *the repo-cyto-GFP* line was generated to drive cytoplasmic GFP expression under the control of the 4.3kb *repo* promoter, which recapitulates the full *repo* expression pattern. [72, 73]

### Hypoxia or hyperoxia rearing conditions

Larvae in a food vial were transferred at 1 day after egg laying (AEL) to a ProOx (model 110, BioSpherix, Lacona, NY) oxygen-controlled chamber. Oxygen or nitrogen was infused into the chamber to a desired concentration (here, 5% or 50%), which was maintained until assay.

### Immunostaining

The NMJ phenotypes were analyzed as previously described [74]. For live tissue preparation, non-fixed larvae were dissected as previously described, before directly incubating larval fillets with anti-horseradish peroxidase (HRP, 1:10) in phosphate buffered saline (PBS) for 10 minutes. Primary antibodies used were against Synapsin (3C11, mouse, 1:100; Developmental Studies Hybridoma Bank, DSHB), HRP-Cy5 (rabbit, 1:100; Jackson ImmunoResearch), Dlg (mouse, 1:100, DSHB), GluRIIA (mouse, 1:100, DSHB), dPAK (rabbit, 1:1000)[75], GluRIII (rabbit, 1:1000)[76], Brp (nc82, mouse, 1:100, DHSB), Sima (guinea pig, 1:1000)[77], Repo (mouse, 1:1000, DSHB), Wg (4D4, mouse, 1:10, DSHB), and Futsch (22C10, mouse, 1:100, DSHB). Secondary antibodies used were anti-rabbit or -mouse 488, Cy3, or Cy5 (1:1000, Jackson ImmunoResearch).

### Image acquisition and processing

Unless specified otherwise, NMJs in muscle 6/7 of A3 segments of wandering third-instar larvae were analyzed. Confocal images were acquired via LSM510 confocal microscopy (Carl Zeiss) using 40x water, 40x water immersion, or 100x oil objectives. All presented images are projections of confocal z-stacks. Numbers of bunch boutons, total boutons, and microtubule loops were counted manually. The immunofluorescence intensities of Wg and HRP, as well as the areas of GFP and HRP signal, were analyzed in ImageJ. Each dot in the bar graph represents the data from a single NMJ of a larva, and 8-10 NMJs from 2-5 independent experiments were pooled for quantification. Embryos were acquired by means of LSM510 confocal microscopy (Carl Zeiss) using a 20x objective, and were analyzed as previously described [36]. Each dot in the bar graph represents data from a single embryo in which fluorescence was measured in at least 35 cells.

### Electrophysiological recordings

Basal transmission properties were analyzed at NMJs of muscle 6/7 in specified segments of wandering third-instar larvae as previously described [78], with some modifications. The larval body wall was dissected in cold calcium-free HL3 solution and recorded in HL3 solution containing 0.4 mM CaCl_2_ at room temperature.

### Crawling behavior

Mid third instar larvae (feeding stage) were placed on black agar plates (2% agar with black food coloring in 25 × 20 cm^2^ dishes) at room temperature for filming. Video recording by a Sony Xperia Z1 camera started after 1 min habituation and lasted for 5 min, and it was analyzed using Ctrax software [79]. The (x, y) positions were used to calculate the crawling distance between two successive frames, and crawling speed was derived by dividing total distance travelled by time. The change in angle of larvae between two frames was divided by time to represent rotational speed. Our forward crawling assay was a modification of a previous study [48]. Larvae were transferred into a tunnel (∼1 mm width) made in 2% black agar. Specimens were video-recorded for 3-10 mins using a Leica S8 APO microscope. Kymographs were constructed using the MultipleKymograph plug-in for ImageJ (NIH). Only forward crawling was counted, and 7-10 steps for each of ten larvae were analyzed for each genotype.

## Acknowledgments

We thank S. Luschnig (Universität Münster), Y. Henry Sun (Academia Sinica), T Leung (National University of Singapore), A. DiAntonio (Washington University), BDSC, Kyoto Stock Center, VDRC, and DSHB for providing reagents; NPAS Electrophysiology Core, Taiwan Fly Core, and Hsiu-Hwa Kao for technical support; as well as Y.-J., Cheng, H. Li, V. Nithianandam and all members of C.-T. Chien’s laboratory for discussion and comments.

## Author contributions

**Conceptualization:** Pei-Yi Chen, Yi-Wei Tsai, Cheng-Ting Chien.

**Formal analysis:** Pei-Yi Chen.

**Funding Acquisition:** Cheng-Ting Chien

**Investigation:** Pei-Yi Chen.

**Methodology:** Pei-Yi Chen, Cheng-Ting Chien.

**Project administration:** Cheng-Ting Chien.

**Resources:** Pei-Yi Chen, Yi-Wei Tsai, Angela Giangrande, Cheng-Ting Chien.

**Supervision:** Cheng-Ting Chien.

**Validation:** Pei-Yi Chen.

**Visualization:** Pei-Yi Chen.

**Writing – original draft:** Pei-Yi Chen.

**Writing – Review & Editing:** Pei-Yi Chen, Yi-Wei Tsai, Angela Giangrande, Cheng-Ting Chien.

## Supporting information

**S1 Fig. Presynaptic and postsynaptic structures in *trh* mutants**

(A) Bar graphs show percentages (mean ± SEM) of satellite boutons to total boutons (*w*^*1118*^, 5.13 ± 1.66%, n = 10; *trh*^*1*^*/+*, 0.66 ± 0.66%, n = 10; *trh*^*1*^*/trh*^*2*^, 2.63 ± 0.97%, n= 10). Statistical significance by Mann-Whitney test is shown as n.s., no significance;, p < 0.5. (B, C, D) Images showing NMJs of muscle 6/7 immunostained for Brp, GluRIII, and HRP (B), GluRIIA, dPAK, and HRP (C), and Dlg and HRP (D) in *w*^*1118*^, *trh*^*1*^*/+*, and *trh*^*1*^*/trh*^*2*^. Scale bars are 10 μm.

**S2 Fig. Tracheal defects in *trh* mutants**

(A) A bright-field view of tracheal dorsal branches of *trh*^*1*^*/+* (left top panel) and *trh*^*1*^*/trh*^*2*^ (right top panel), and tracheal dorsal trunks of *trh*^*1*^*/trh*^*2*^ (bottom panels). Numbers denote terminal branches, and arrows indicate a tracheal break (bottom left) and a tracheal tangle (bottom right). Scale bar represents 50 μm. (B) Bar graph shows mean ± SEM of dorsal terminal branches (*trh*^*1*^*/+*, 5.70 ± 0.15, n = 10; *trh*^*1*^*/trh*^*2*^, 7.55 ± 0.28, n = 11). (C) Bar graphs show mean ± SEM of the GFP immunofluorescence intensity (*w*^*1118*^ at 21% O_2_, 20.33 ± 7.54, n = 6; *w*^*1118*^ at 5% O_2_, 79.31 ± 17.98, n = 6; *trh*^*1*^*/+* at 21% O_2_, 13.29 ± 3.64, n = 6; *trh*^*1*^*/trh*^*2*^ at 21% O_2_, 108.2 ± 14.70, n = 6). (D) Bar graphs show mean ± SEM of RFP immunofluorescence intensity (*w*^*1118*^ at 21% O_2_, 102.00 ± 15.55, n = 6; *w*^*1118*^ at 5% O_2_, 109.60 ± 13.12, n = 6; *trh*^*1*^*/+* at 21% O_2_, 143.00 ± 15.67, n = 6; *trh*^*1*^*/trh*^*2*^ at 21% O_2_, 151.50 ± 16.77, n = 6). Statistical significance by Mann-Whitney test is shown as n.s., no significance; **, p < 0.01; ***, p < 0.001.

**S3 Fig. Efficiency of *sima-RNAi* knockdown and Sima upregulation**

(A) Assay for *sima* mRNA levels in *da-GAL4* (control) or *da-GAL4/sima-RNAi* shows a reduction upon *sima-RNAi* knockdown. The *Rpl19* mRNA levels are comparable in both genotypes. (B) Images showing central nerve cord immunostained for Sima (green) and Repo (red) in *w*^*1118*^ at 21% O_2_, *w*^*1118*^ at 5% O_2_, and *trh*^*1*^*/trh*^*2*^ at 21% O_2._ Arrows indicate glia with high Sima protein levels. Scale bar represents 20 μm.

**S4 Fig. Microtubule loops and glial invasion of *trh* NMJs**

(A) Images showing NMJs of muscle 6/7 immunostained for HRP (red) and Futsch (green) in *trh*^*1*^*/+* and *trh*^*1*^*/ trh*^*2*^. White arrows indicate microtubule loops. Dashed squares are enlarged in right panels. Scale bar represents 10 μm. (B) Bar graphs show averages (mean ± SEM) of Futsch loops (*trh*^*1*^*/+*, 2.92 ± 0.29, n = 12; *trh*^*1*^*/ trh*^*2*^, 9.56 ± 1.11, n = 9). (C) 3D images of live tissues showing NMJs of muscle 6/7 carrying *repo-cyto-GFP* (green) labeled for HRP (red) in *trh*^*1*^/+ and *trh*^*1*^*/trh*^*2*^ by using Imaris. Synaptic overlap by glia is represented in yellow. Scale bar represents 10 μm. (D) Bar graph shows percentages (mean ± SEM) of synaptic areas overlapping with glia in *trh*^*1*^*/+*, 6.2 ± 0.99%, n = 10; *trh*^*1*^*/trh*^*2*^, 16.7 ± 3.66, n = 10. Statistical significance by Mann-Whitney test is shown as *, p < 0.05 or ***, p < 0.001.

**S1 Movie. Forward crawling of *w****^1118^*****

**S2 Movie. Forward crawling of *trh****^1^****/trh****^2^*****

