## Supplementary figures and images for "Glial response to hypoxia in *trachealess* mutants induces synapse remodeling"

### Supporting information

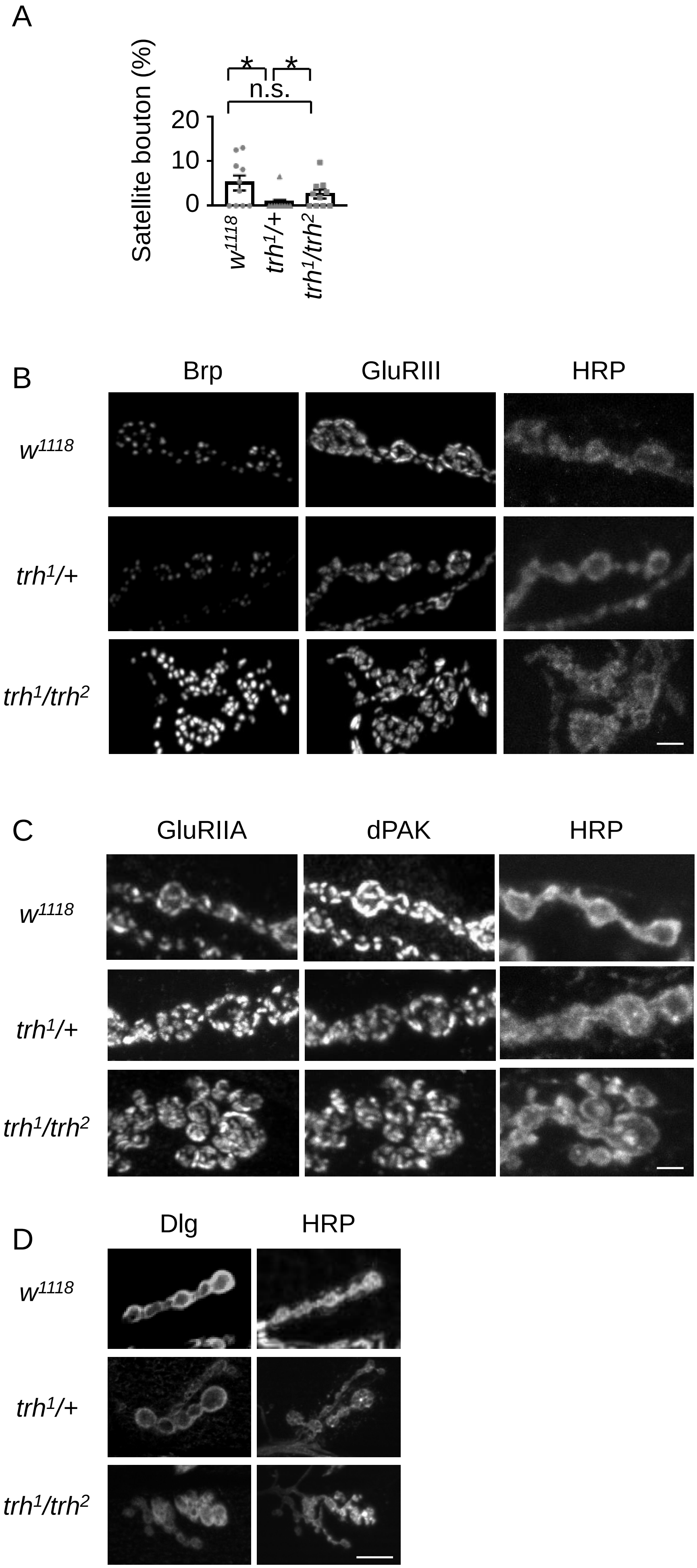

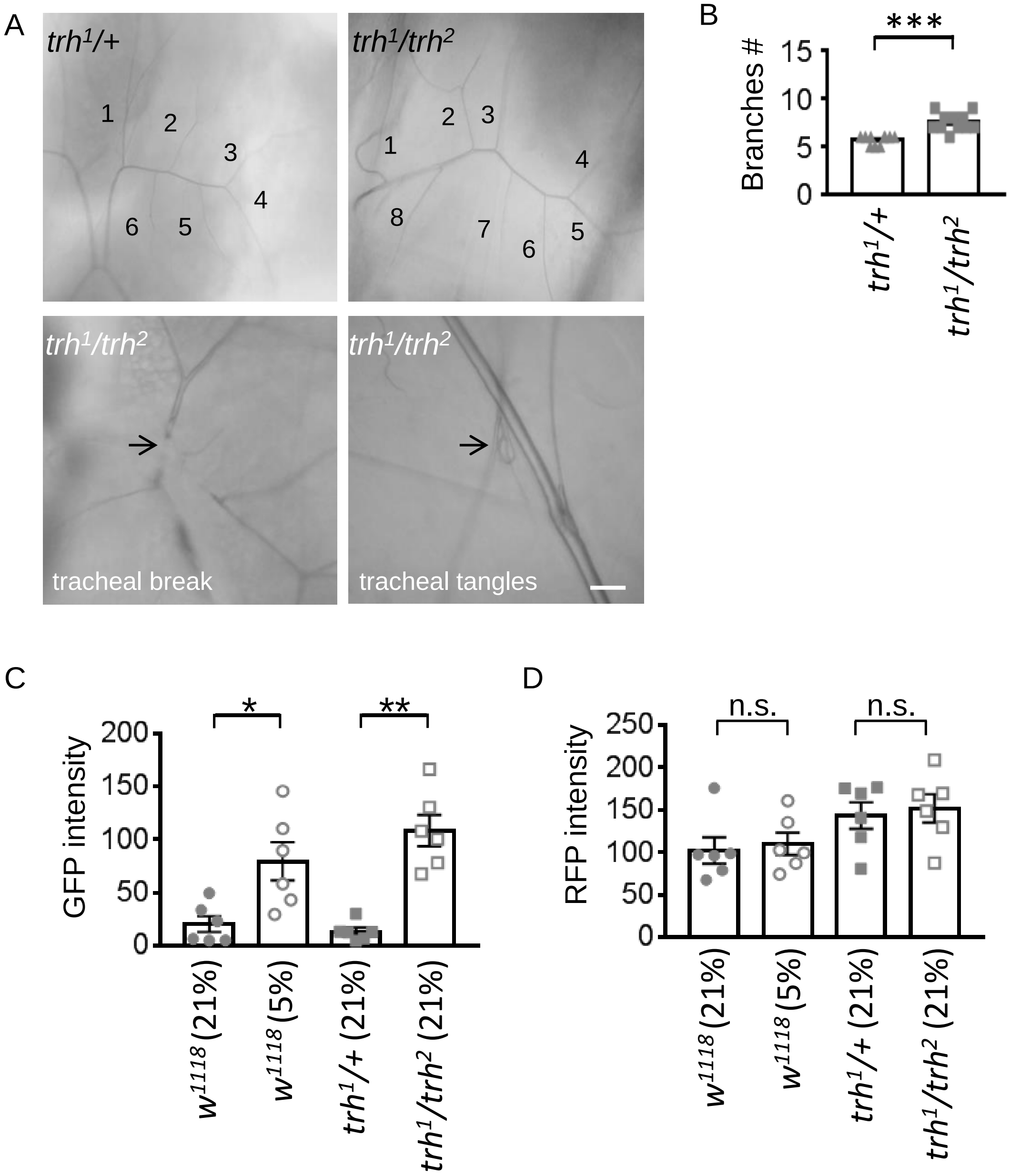

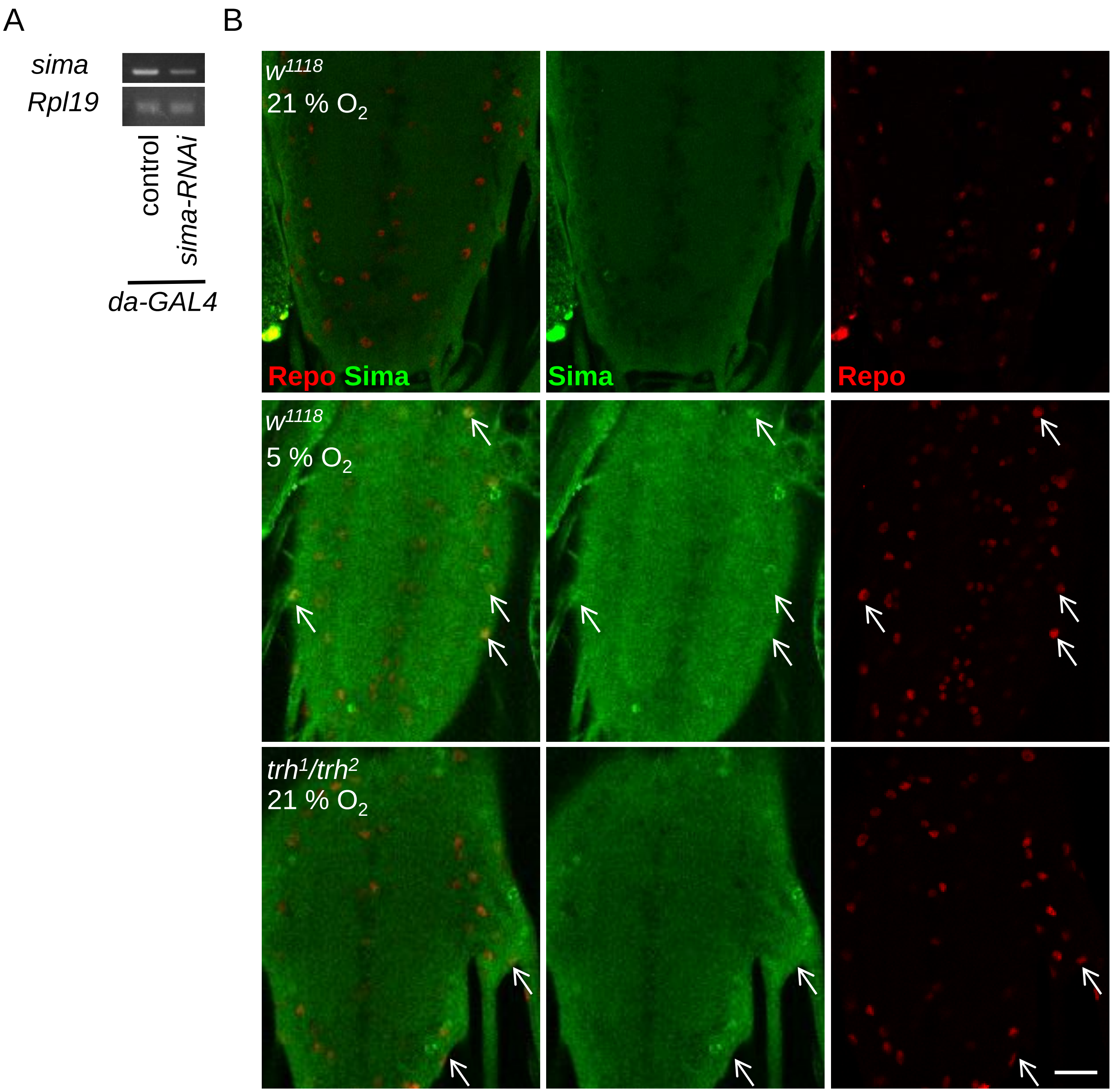

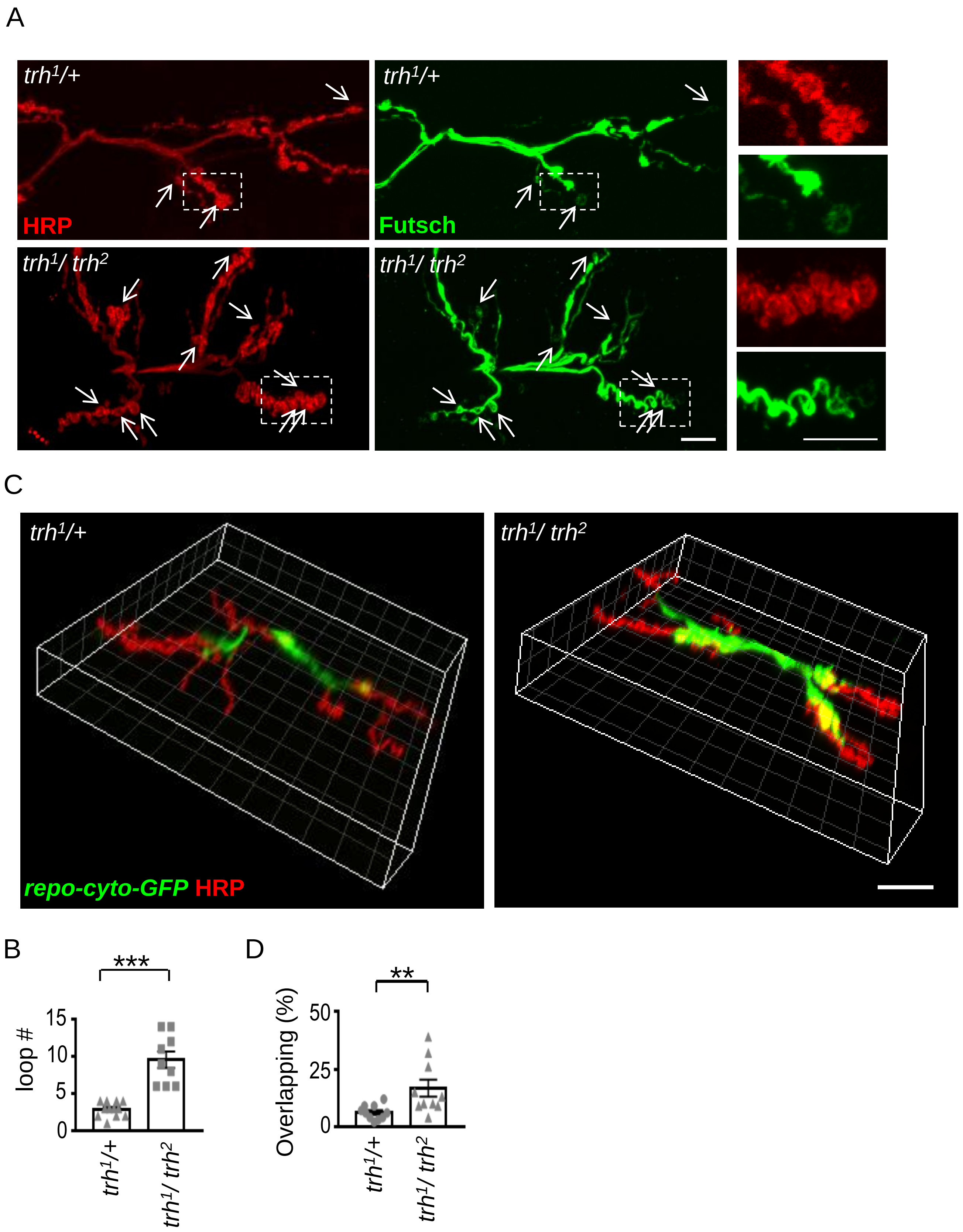
